# Biochemical and Immuno-Histochemical Localization of Type IIA Procollagen in Annulus Fibrosus of Mature Bovine Intervertebral Disc

**DOI:** 10.1101/2021.03.26.437279

**Authors:** Audrey McAlinden, David M. Hudson, Aysel A. Fernandes, Soumya Ravindran, Russell J. Fernandes

**Affiliations:** Department of Orthopaedic Surgery, Washington University School of Medicine, St Louis, MO, USA; Department of Cell Biology & Physiology, Washington University School of Medicine, St Louis, MO, USA; Shriners Hospitals for Children- St Louis, MO, USA; Department of Orthopaedic & Sports Medicine, University of Washington, Seattle, WA, USA

**Keywords:** Type IIA procollagen, alternative splicing, type II collagen, nucleus pulposus, annulus fibrosus, collagen fibrils; extracellular matrix, intervertebral disc, cartilage, mass-spectrometry, post-translational modifications, Prolyl-3-hydroxylase 2; 3-hydroxyproline

## Abstract

For next generation tissue-engineered constructs and regenerative medicine to succeed clinically, the basic biology and extracellular matrix composition of tissues that these repair techniques seek to restore have to be fully determined. Using the latest reagents coupled with tried and tested methodologies, we continue to uncover previously undetected structural proteins in mature intervertebral disc. In this study we show that the “embryonic” type IIA procollagen isoform (containing a cysteine-rich amino propeptide) was biochemically detectable in the annulus fibrosus of both calf and mature steer intervertebral discs, but not in the nucleus pulposus where the type IIB isoform was predominantly localized. Specifically, the triple-helical type IIA procollagen isoform immunolocalized in the outer margins of the inner annulus fibrosus. Triple helical processed type II collagen exclusively localized within the inter-lamellae regions and with type IIA procollagen in the intra-lamellae regions. Mass spectrometry of the α1 (II) collagen chains from the region where type IIA procollagen localized showed high 3-hydroxylation of Proline-944, a post-translational modification that is correlated with thin collagen fibrils as in the nucleus pulposus. The findings implicate small diameter fibrils of type IIA procollagen in select regions of the annulus fibrosus where it likely contributes to the organization of collagen bundles and structural properties within the type I-type II collagen transition zone.

## INTRODUCTION

In the adult spine, the function of carrying and transmitting mechanical compressive load, as well as flexibility that allows bending and twisting, rests on the intervertebral discs (IVD) that join adjacent vertebral bodies [1–3]. Anatomically, the mature disc consists of three distinct but integral regions, (i) the soft inner nucleus pulposus (NP) in the center of the disc, (ii) the robust annulus fibrosus (AF), a specialized fibrocartilage that laterally encloses the NP, and (iii) the cartilaginous end plate that vertically encloses the NP [4, 5].

The main structural macromolecular components of all the regions are collagens (types I and II) and proteoglycans (aggrecan, biglycan, decorin, fibromodulin) [6–13]. Types III, IX, XI and VI collagens have also been found in minor amounts [14–17]. The collagens polymerize into fiber bundles in the AF lamellae that are concentrically formed around the NP [18]. Although the precise function of all types of collagens in the IVD is still not clear, it is established that type I and type II collagens are responsible for the load bearing properties [7]. There is a decreasing gradient of type II collagen from the NP to the outermost lamellae of the AF and conversely a decreasing type I collagen gradient from the outer AF to the NP where this collagen is not present [6]. Type II collagen is essential for the removal of the notochord and the formation of IVD [19]. The other fibrillar structural proteins, elastin and fibrillin, have also been detected within the lamellae of the AF of the IVD [20, 21]. Fibrillin is present in radial cross-bridging elements within AF lamellae of sheep discs, bovine tails, and adult human discs [22] that appear to be a consequence of vascular regression during development [23, 24].

The inter-lamellar architecture within the AF is of intense biochemical interest as it has been shown that transverse microscale deformation occurs as a result of interlamellar skewing and not sliding between annular layers [25]. Immunohistochemical imaging of normal bovine and human AF suggests that interlamellar connections involve a variety of molecular interactions including collagen, elastin and fibrillin-1 [22, 26] [24]. Collagen and fibrillin, rather than elastin, contribute to the rigid connection between the lamellae rat IVD [25]. Scanning electron microscopy of rat AF also revealed large diameter collagen fibers that contained many small diameter fibrils. This study also revealed an interesting micro-ultrastructural feature where the collagen in the AF formed distinct small tubules of inner diameters 1-2 μm that appeared to run along the length of the collagen fibers in the AF [27]. Similar tubules were also observed in the radial zones of articular cartilage [28]. The composition of this tubular network is still unclear, prompting further investigation of previously unidentified collagen types in this fibrocartilaginous tissue.

Of interest to this study are the alternatively-spliced isoforms of type II collagen. Like all fibrillar collagens, type II collagen is generated first as precursor procollagen containing a triple helical domain flanked by a 5’-amino and a 3’-carboxy propeptide. Alternative splicing of the type II procollagen gene (*Col2a1*) has been well described during cartilage development whereby the IIA isoform (generated by precursor chondrocytes) is formed by inclusion of exon 2 in the amino propeptide while the IIB isoform (synthesized by differentiated chondrocytes) is generated by exclusion of exon 2 [29–32]. This developmentally-regulated alternative splicing event has also been detected during embryonic development of the human IVD, particularly in the inner annulus and NP [33]. Interestingly, previous studies have reported apparent re-expression and matrix deposition of this “embryonic” IIA procollagen isoform in osteoarthritic cartilage [34] as well as within degenerated adult human intervertebral disc tissue [35]. We have also shown that transgenic mice engineered to synthesize only the IIA procollagen isoform developed normally and deposited this collagen isoform into cartilage extracellular matrix (ECM) during adulthood [36, 37]. Given these observations that the IIA procollagen can be deposited in adult connective tissues, we explored the possibility that this “embryonic” isoform may be present within the annulus fibrosis of healthy mature bovine and calf intervertebral disc tissue.

We show here that the type IIA procollagen isoform is biochemically detectable in the AF of both mature steer and calf IVD and that it immunolocalizes to the outer margins of the inner AF. Furthermore, by mass spectrometry, we determined that the α1 (II) collagen chains from the defined area containing the type IIA procollagen isoform shows specific post-translational hydroxylation modifications often found associated with thin collagen fibrils. These novel findings demonstrate the presence of type IIA procollagen at the type I-II collagen transition zones in the annulus fibrosus of post-natal intervertebral disc tissue, thereby suggesting an important and previously unrecognized role for this isoform in tissue ultrastructure or in maintaining tissue homeostasis.

## METHODS

### Tissue Specimens

Intervertebral discs from healthy adult steer tail IVD (4 year old) and calf (3 month old) spines were obtained fresh from the local butcher/abattoir and used immediately for collagen analyses or frozen at −80°C for immunohistochemistry. Fetal bovine spinal columns (2-3 month gestation) were obtained from Sierra (Whittier, CA). Lumbar discs L3 and L4 were dissected and frozen at −80°C for isolation of collagen.

### Collagen extraction and analysis

Clean gelatinous white NP was first dissected from the adult IVDs. Next, a region of the inner annulus adjacent to the NP was dissected and designated as the Inter-zone. Moving radially outward, the region of the inner annulus (IA) and the outer annulus (OA) was dissected and designated as the annulus fibrosus (AF). The outermost tough AF was discarded (see Figure 1A, B). Disc tissues were rinsed with PBS and proteoglycans, non-cross-linked and/or newly synthesized collagen were extracted with 50mM Tris buffered 4M Guanidinium HCl (GuHCl) for 24 h at 4°C [38, 39]. Following thorough dialysis in distilled water, the extracts were lyophilized. Type II collagen in the residue was solubilized with pepsin (0.5mg/ml) in 0.5M acetic acid (pH 3) for 18 hours at 4°C [38, 40] and used for mass specrometry. Mouse type IIA procollagen from 4M GuHCl extracts of rib cartilage from the type IIA procollagen knock-in mouse (ki/ki) [36, 37] was used as a control. Control type IIB collagen was extracted from the Swarm rat chondrosarcoma RCS-LTC cell line that we have shown to synthesize only the type IIB procollagen isoform [41].

**Figure 1.**
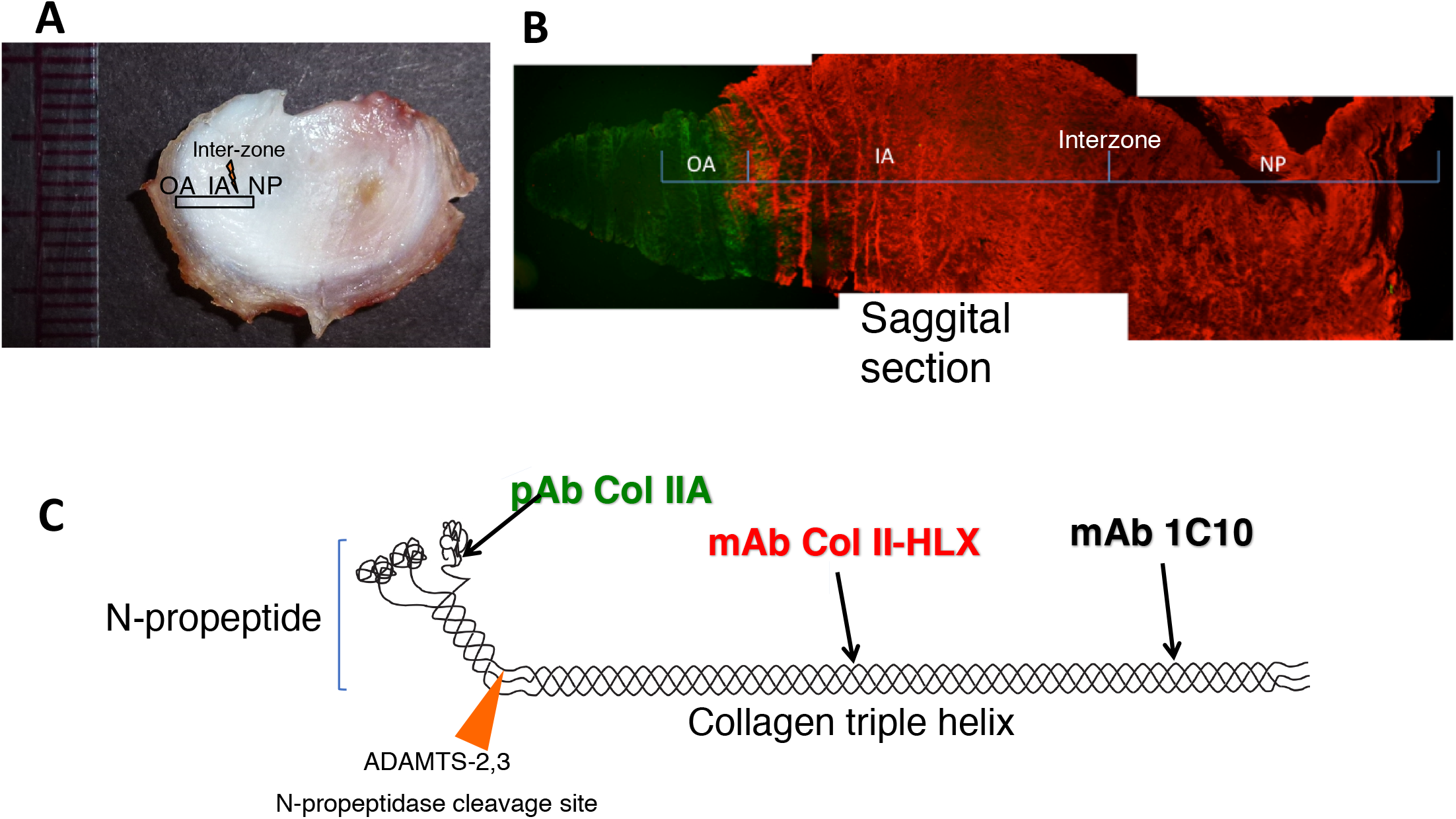
Intervertebral disc tissue sampling and antibodies used to detect type IIA procollagen. A) Bovine steer tail whole intervertebral disc showing area dissected for immunofluorescence staining. OA, outer annulus, IA, inner annulus, the interzone and NP, nucleus pulposus. B) Composite image of sagittal sections showing regions of the disc containing OA, IA, the inter-zone and NP. Note the green fluorescence staining in the OA region indicating type IIA procollagen and red fluorescence staining indicating the triple helical domain of type II collagen. C) Cartoon showing the structure of type IIA N-procollagen containing the amino propeptide (N-propeptide). Location of epitopes recognized by each antibody are also highlighted: Col IIA antibody binds to the exon 2 encoded cysteine-rich (CR) domain in the N-propeptide; Col II-HLX antibody binds to the triple helical domain of type II collagen; 1C10 antibody recognizes a region of the triple helical domain closer to the C-terminal end. Col IIA and Col II-HLX antibodies were used for immunofluorescence staining and Col IIA and 1C10 antibody was used for Western blots. The region of the ADAMTS-2 or 3 N-propeptidase cleavage site is also shown. These enzymes process type II N-procollagen to form mature type II collagen.

For smaller calf IVDs, the clear transparent NP was removed and the entire annulus fibrosus was dissected and kept intact. Proteoglycans, non-cross-linked collagen and newly synthesized collagen from the AF was extracted with 4M GuHCl as described above. Following 4M GuHCl extraction, residual proteoglycans and procollagens in the calf AF residue were extracted further by treating first with 0.125 unit/ml chondroitinase ABC (EC4.2.2.4) (Seikagaku Kogyo Co., Tokyo, Japan) in 4 ml of 0.05 M-Tris/ HC1/0.06M-sodium acetate buffer, pH8.0, containing 2mM-phenylmethanesulphonyl fluoride, 2mM-EDTA, 5 mM-benzamidine and 10mM-N-ethylmaleimide at 22 °C (room temperature) for 24 h and then with 0.5M NaCl in 50mM Tris-HCl, pH 7.5 for 25 hours at 4 °C [16, 42]. These chondroitinase ABC extracts were then dialyzed against distilled water and lyophilized. The 0.5M NaCl extracts were dialyzed against 0.1M acetic acid, centrifuged and the supernatant (acid soluble) and residue (acid insoluble) were lyophilized separately.

The NP from calf disc was highly hydrated and near transparent. Non-cross-linked collagen from calf discs was easily extracted with chondroitinase ABC leaving a highly cross-linked mature collagen network in the residue that was not analyzed further. Aliquots of all lyophilized material were dissolved in Laemmli sample buffer for electrophoresis.

### Western blots

Intact and processed type II collagen chains in tissue extracts were resolved by SDS-PAGE (6% gel), transferred to PVDF membrane and probed with monoclonal antibody to type II collagen (1C10) which recognizes a domain in the triple helical region [43–45] and a polyclonal antibody to type IIA procollagen which recognizes the exon 2-encoded domain in the N-propeptide of type IIA procollagen (Col IIA) [36, 46] (Figure 1C). Intact and processed type IIA/IIB collagen chains were identified by molecular weight and migration pattern as we have done before [37, 41]. For type IIA procollagen control we used type II collagen extracted from the rib cartilage of a transgenic mouse (ki/ki) engineered to synthesize only the IIA procollagen isoform [37].

### Immunofluorescence Staining

Cryo-sections (8-10 μm, sagittal orientation) of the AF and NP were cut and incubated with 1% hyaluronidase (Sigma) for 30 min at 37°C [20]. Sections were rinsed with PBS and blocked with 10% goat serum for 1 h at room temperature and then incubated overnight at 4°C with the rabbit polyclonal anti-type IIA antibody (Col IIA) that recognizes the exon 2-encoded cysteine-rich domain within the amino-propeptide of type II procollagen [33, 35, 46], a rat polyclonal antibody against the triple helical domain of type II collagen (Col II-HLX) [47]. Each antibody was diluted 1/400, and 1/100 respectively in 2% goat serum. Following 1x PBS washes, sections were incubated with species-specific secondary antibodies (1/250 dilution) that were conjugated to Alexa fluorescent dyes (Invitrogen: goat anti-rabbit Alexa 488; goat anti-rat Alexa 594) for 1 h at room temperature. DAPI mounting medium was applied following three rinses in PBS and stained sections were cover-slipped. A Nikon Eclipse E800 fluorescence microscope was used to view the fluorescent images. The FITC and TRITC band pass filter sets were used to view sections labeled with Alexa 488 (green) and 594(red) dyes, respectively and the DAPI filter set was used for viewing cell nuclei [37]. Figure 1C shows the specificity of antibodies to the type IIA procollagen molecule.

### Mass Spectrometry

Regions of bovine steer IVDs shown, by immunohistochemistry, to be enriched in both type I and type IIA collagens were carefully dissected. These regions encompassed the most distal part of the inner annulus (relative to the NP) together with the outer annulus fibrosis. Type II collagen from adult NP, adult inner AF, adult outer AF, fetal NP, fetal AF, adult bovine articular cartilage, fetal bovine rib, spinal process cartilage and adult bovine vitreous was solubilized with pepsin as described above and analyzed by mass spectrometry. To compare 3-Hydroxyproline (3Hyp) occupancies in type II collagen before birth and in adult, fetal rather than calf tissue was analyzed. Pepsin solubilized collagen chains were resolved by SDS-PAGE under reducing conditions and identified by Coomassie Blue staining. Individual collagen a-chains were excised and subjected to in-gel trypsin digestion [48, 49]. Electrospray MS was performed on the tryptic peptides using an LTQ XL linear quadrapole ion-trap mass spectrometer equipped with in-line liquid chromatography (ThermoFisher Scientific) using a C8 capillary column (300 x150 mm; Grace Vydac 208MS5.315) as we have described before [50, 51]. Thermo Xcalibur software and Proteome discoverer software (ThermoFisher Scientific) were used for peptide identification using the NCBI protein database. Collagenous peptides not found by by the software had to be identified manually by calculating the possible MS/MS ions and matching these to the actual MS/MS. The percentage 3Hyp at a particular site was determined from the abundance of 3Hyp-containing peptide ions as a fraction of the sum of both 3Hyp and Pro versions of the same tryptic peptide [36].

## RESULTS

### Biochemical identification of type IIA procollagen in bovine IVD

Western blot analysis showed the presence of type II procollagen chains containing the IIA amino propeptide and devoid of the carboxy propeptide [pN-α1(IIA)] as a single band (~150 KDa) in the 4M GuHCl extract of adult steer AF (lane 3) but not in the NP and the AF/NP inter-zone (lanes 4-5) (Figure 2A). A robust signal was observed for control pN-α1(IIA) chain (lane 1), from the transgenic mouse (ki/ki) engineered to synthesize only type IIA isoform as we have shown previously [37]. As expected no reactivity was observed for the type IIB procollagen extracted from the RCS-LTC cell line (lane 2) [41].

**Figure 2.**
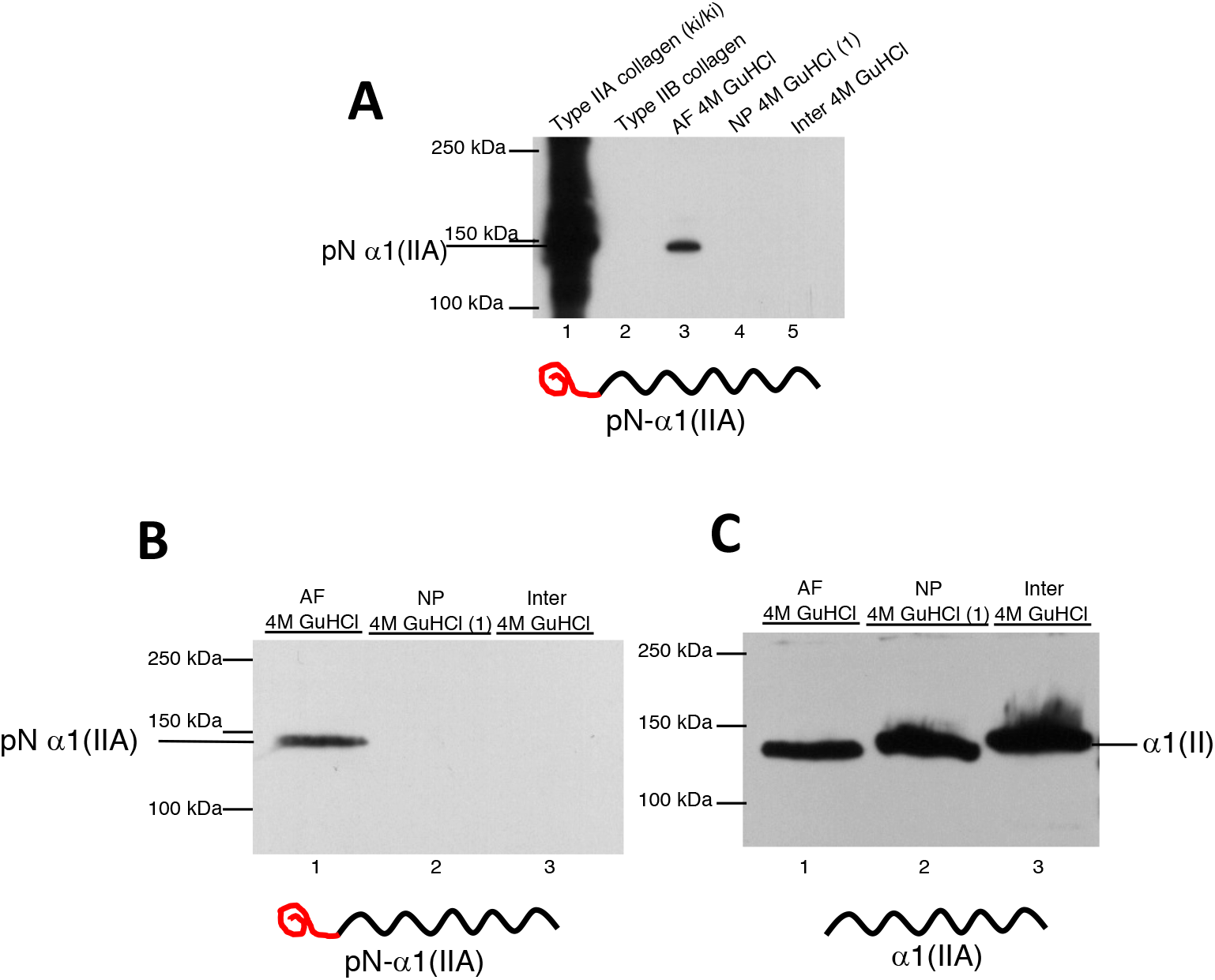
Detection of type IIA procollagen in steer annulus fibrosus. A) Western blot analysis using established type IIA collagen antibodies showed the presence of the pN-α1(IIA) chain as a single band, in the 4M GuHCl extract of AF (lane 3) but not in the samples of NP and the AF interzone (lanes 4-5). A robust signal was observed for control type IIA procollagen (lane 1) and no signal for type IIB collagen as expected (lane 2). Note the line drawing under the western blot showing that pN-α1(IIA) comprises the type II collagen triple helical domain (black) bound to the IIA amino propeptide containing the exon 2 encoded cysteine-rich domain (red). B) Probing a different blot with wider lanes and higher protein loads of IVD extracts, confirmed the presence of pN-α1(IIA) chain only in the AF. C) Probing an identical blot to B, the type II collagen antibody 1C10, revealed the abundant presence of processed α1(II) collagen chains (containing only the helical domain) in NP, AF and the interzone. Note the slower migration of the pN-α1(IIA) chain band in (A) and (B) as compared to the fully processed α1(II)chain band shown here in (C).

Since we had expected to detect type IIA procollagen chains in healthy bovine NP, but not in bovine AF, the data in Fig. 2A prompted us to probe another blot of IVD extracts containing higher concentrations of protein extracts per lane. This western blot (Figure 2B) confirmed a solitary band in the same molecular weight range (~150 KDa) as the type IIA procollagen chains in the AF (Figure 2A). In an identical blot, the monoclonal antibody to the triple helical region of α1(IIA/B) chain (1C10), revealed abundant fully processed α1(II) collagen chains in NP, AF and the NP/AF inter-zone extracts (Figure 2C). Note the slower migration of the pN-α1(IIA) band in Figure 2B, lane 1 when compared to the processed α1(II) band in Figure 2C, lanes 1-3).

The unexpected presence of type IIA procollagen in adult steer AF but not in NP prompted us to examine younger calf intervertebral disc tissues. We used classical reagents known to extract non-cross-linked type V procollagen, type VI and type II procollagen based on their interactions with other molecules in the extracellular matrix. [16, 37, 42]. As shown in Figure 3A, pN-α1(IIA) chains were not detected in the chondroitinase ABC enzyme digests of both AF (lane 1) and NP (lane 5). In the chaotropic 4M GuHCl extracts of AF (lane 2) pN-α1(IIA) chains were identified as a single band similarly detected in the adult AF (Figure 2). The high salt 0.5M NaCl extracts showed strongly reactive bands in the range of pN-α1 (IIA) chains and pro-α1(IIA) chains in the acid-soluble fraction (lane 3). The acid-insoluble fraction showed a faint reactivity to only pro-α1(IIA) chains (lane 4). A probe of an identical blot with the type II collagen monoclonal antibody 1C10 with an epitope in the triple helical region of α1(IIA/B) chain, (Figure 3B) revealed abundant and pN-α(IIB) collagen chains in the chondroitinase ABC digest as well as both pro α1(IIB) and pN-α(IIB) in the other extracts. This antibody was also able to identify, although faintly, pN-α1(IIA) and pro-α1(IIA) bands (Fig 3B, lane 3, 4, white asterisks) in the 0.5M NaCl extracts. We have published on a similar pattern of native and unprocessed collagen for type IIA/type IIB collagen in epiphyseal cartilage from one week old wild type, homozygous and heterozygous type IIA procollagen knock-in mice [37]. The presence of type II collagen processing intermediates is indictive of collagen synthetic activity which is reflected in calf disc. In adult disc where synthetic activity is low there is a near absence of unprocessed type II collagen chains (Figure 2). These results, using chaotropic, salt and enzymatic extraction agents, indicate differences in type IIA collagen molecular assembly and/or differential interaction of this collagen with other collagens or matrix components.

**Figure 3.**
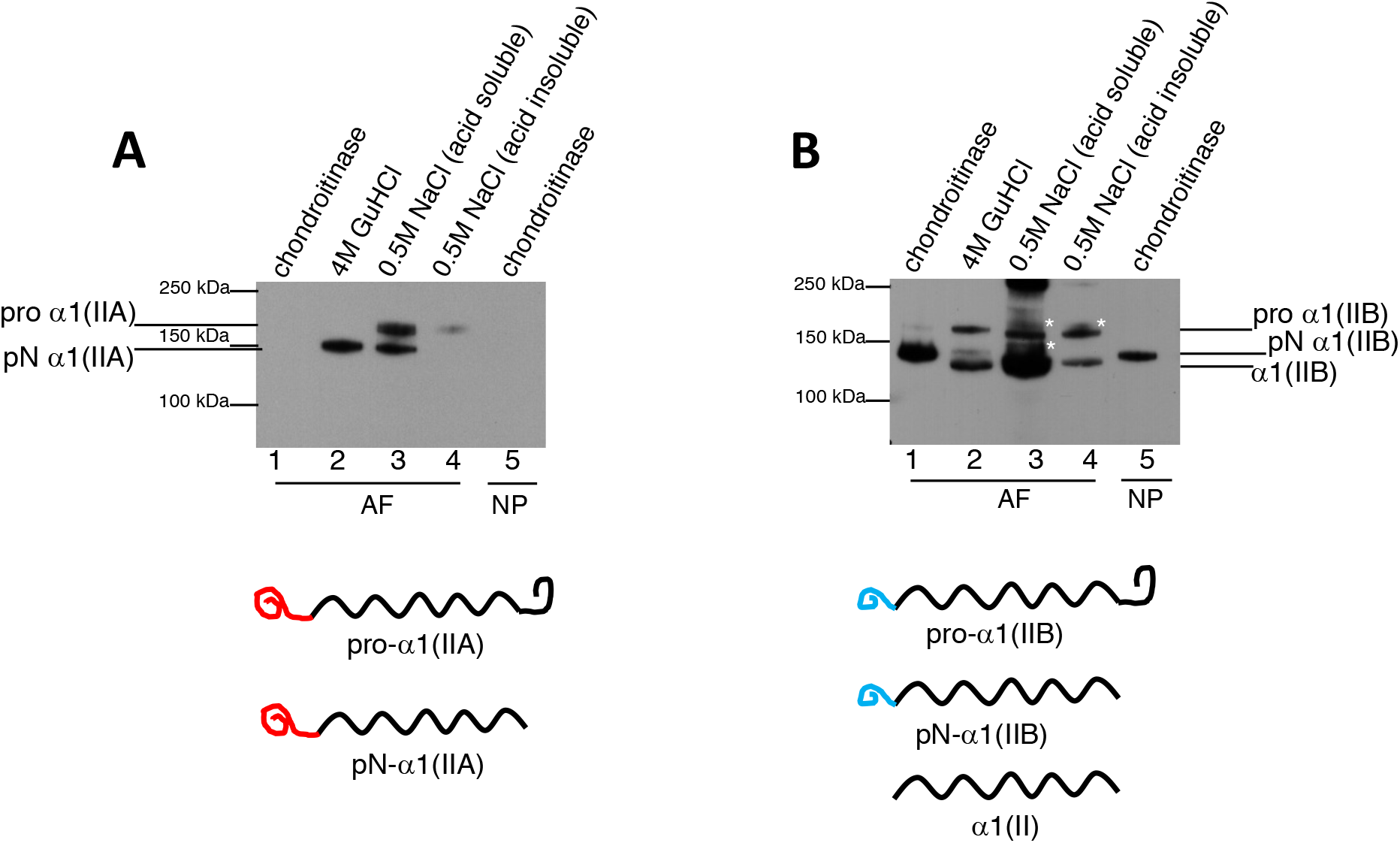
Detection of type IIA procollagen in calf annulus fibrosus. A) In western blots, type IIA collagen antibodies (Col IIA) also identified the pN-α1(IIA) chain and the slower moving, pro-α1(IIA) collagen chain in the 0.5M NaCl extract (acid soluble) of calf AF (lane 3), with faint reactivity to the pro-α1(IIA) chain in the acid insoluble fraction (lane 4). No pN-α1(IIA) chains were detected in the chondroitinase ABC digests of both AF and NP (lane 1, 5). In 4M GuHCl extracts, pN-α1(IIA) chains were identified as a single band (as expected from Figure 2). B) The type II collagen antibody (1C10) revealed an abundance of, pN-α1(IIB) collagen chains in the chondroitinase digest (lanes 1, and 5) as well as the 4M Gu HCl extract. Pro-α1(IIB) chains were also observed in the 4M GuHCl and the 0.5 M NaCl extracts (lanes 2-4). The faint bands in the 0.5M NaCl soluble and insoluble extracts marked with a white asterisk (*) are in the same molecular weight range as pN-α1(IIA) and pro-α1(IIA) chain bands. These extraction results indicate differences in type IIA collagen molecular assembly or differential interaction of this collagen with other matrix components. Note the line diagrams underneath the western blots shown in (A) and (B) indicating the (pro)collagen forms identified in the labeled protein bands. Red line (IIA amino propeptide containing exon 2-encoded cysteine-rich domain), blue line (IIB amino propeptide devoid of the cysteine-rich domain), black line (helical domain), black loop (carboxy propeptide).

### Immunolocalization of type IIA procollagen in the mature IVD

To precisely localize type IIA procollagen and triple helical type II collagen protein, we used the type IIA propeptide antibody (Col IIA) and the antibody that recognizes the triple helical domain of type II collagen (Col II-HLX) in sagittal sections of steer IVD. As seen in Figure 4 (A-D) upper panel, the α1(IIA) N-propeptide of type IIA collagen (Col IIA, green) was clearly detected in the outer annulus (Fig. 4A) and within the intra-lamellar regions in the outer area of the inner annulus (Fig. 4B, white arrows) but was not detected in the interzone closer to the NP (Fig. 4C) or in the NP (Fig. 4D). Immuno-localization of triple-helical type II collagen (Col II-HLX, red) clearly shows no triple helical type II collagen was detected in the outer annulus (Fig. 4E) but was present in the inner annulus adjoining the outer annulus (Fig. 4F). In this region, triple helical type II collagen seemed to be concentrated within the inter-lamellar regions (Fig. 4F, vertical red tracks indicated by orange arrows), but was also detected in the intra-lamellar regions. Abundant helical type II collagen in the extracellular matrix of the interzone closer to the NP (Fig. 4G) and in the NP (Fig. 4H). When both the signals were merged [Figure 4 (I-L) lower panel], it was clear that the Col II-HLX (red) signal was more localized to the inter-lamellar regions of the inner annulus (J, vertical red tracks, orange arrows) and both Col II-HLX and Col IIA (green) signals co-localized in intra-lamellar regions and was seen as yellow/lighter green (J, white arrows). This indicated triple helical type IIA procollagen in the intra-lamellar regions. Col II-HLX did not co-localize with Col IIA (Fig 4I) in the outermost regions of annulus.

**Figure 4.**
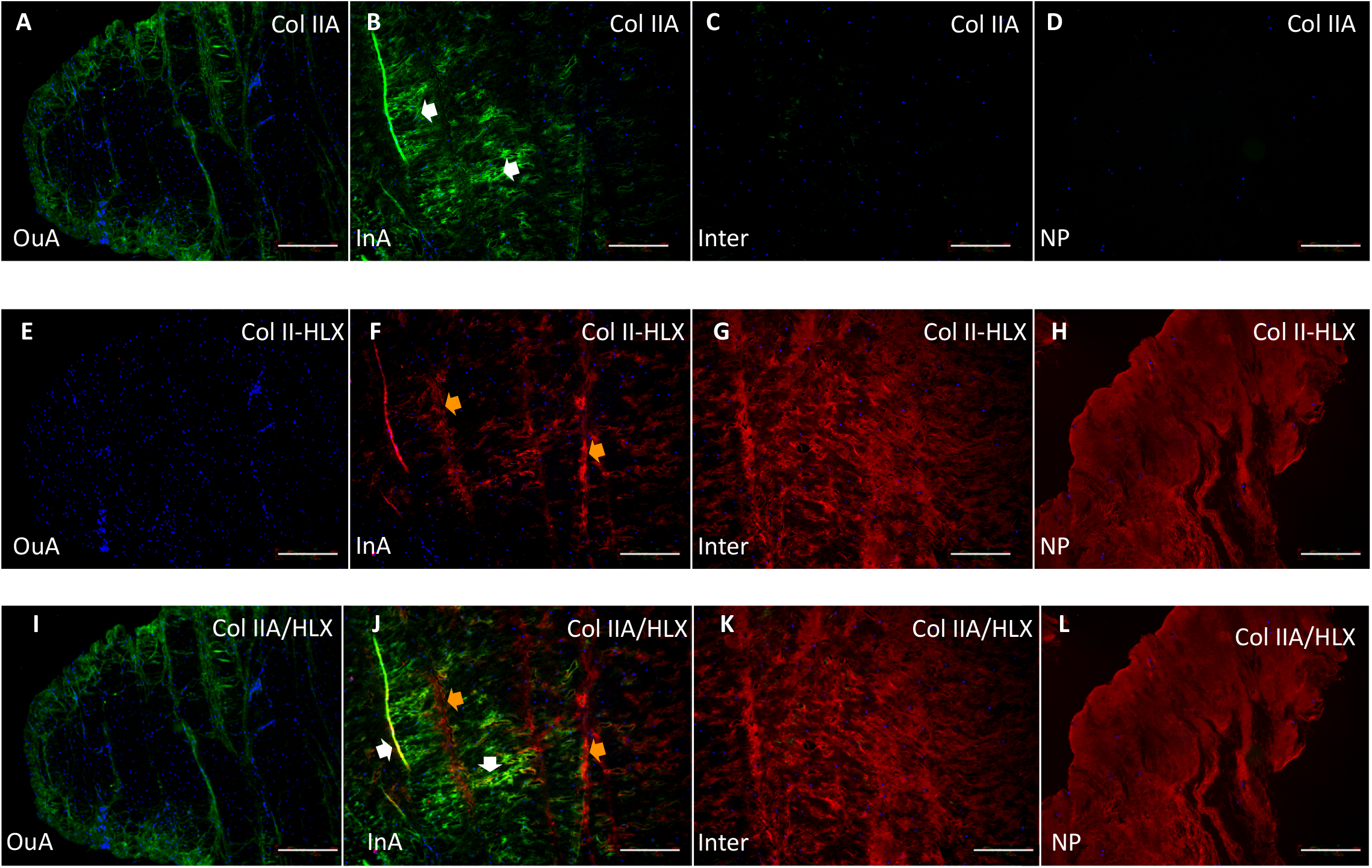
Fluorescence immunolocalization of type IIA procollagen in steer IVD. Sagittal sections of steer intervertebral disc probed with type II collagen antibody (Col II-HLX, red), and Col IIA, green) antibodies. Panels A-D, Col IIA staining of outer annulus (OuA), inner annulus (InA), interzone, region between InA and NP (Inter), nucleus pulposus (NP). Panels E-H, Col II-HLX staining of OuA, InA, Inter, and NP. Panels I-L, merged Col IIA and Col II-HLX staining in OuA, InA, Inter, and NP. Col IIA signal (green) was detected in the intra-lamellar regions of the inner annulus (B, white arrow) and the OuA (A). In InA, Col II-HLX (red) seemed to be concentrated in the inter-lammelar regions (F, orange arrows). When both the signals were merged, it was clear that both Col II-HLX (red) and Col IIA (green) signals localized together (J, light green/yellow signal, white arrows). In the inter-lamellar regions of the inner AF (J, orange arrow) only the Col II-HLX (red) signal was seen. Punctate blue DAPI stained nuclei were clearly seen in all sections. Scale bars = 100μ

This co-localization is better visualized in Figure 5. Type IIA procollagen (green) and helical type II collagen (red) were both clearly localized in the extracellular matrix (Fig, 5B), and were merged in areas where co-localized (yellow). In Fig. 5F, a higher magnification of the boxed area in B shows co-loclization in even more detail within the intra-lamellar region of the inner annulus and closer to the outer annulus (white arrows). This section however, also showed regions where type IIA propeptide and triple helical type II collagen (F, white asterix) were not merged and distinctly separate in localization. There was no co-localization of type IIA procollagen and helical type II collagen in any of the other regions even under high magnifications (Fig. 5, panels A, C D,E G, H)

**Figure 5.**
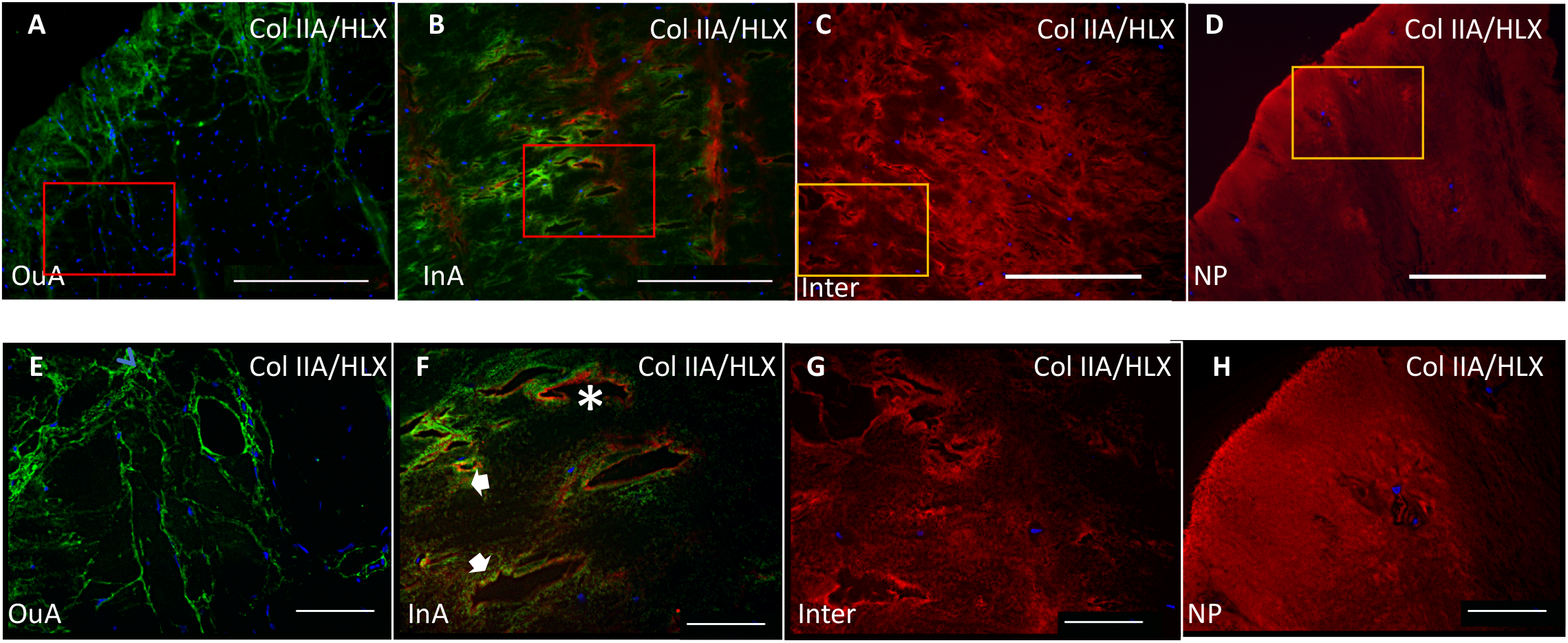
Co-localization of type IIA procollagen and triple helical type II collagen <u>Panels A - D show</u> higher magnifications of the bottom panel of Figure 4. <u>Panels E, F, G, H are higher magnification images of the boxed regions in A, B, C, D, respectively</u>. Type IIA collagen (green, A, B, E, F,) and helical type II collagen (red, B, C, D, F, G, H) were both clearly localized in the extracellular matrix, and when merged, were co-localized (light green/yellow) in some areas of the inner annulus (B) and seen clearly in a higher magnification of the boxed area (F, white arrows). Regions where type IIA collagen and type II collagen were not merged and distinctly separate in localization were also seen (F (white asterix). Nuclei are stained blue and can be seen in all regions of the IVD. Bar = 100μ in A, B, C, D. and 20μ in E, F, G, H.

### Differences in post-translational Proline 3-hydroxylation in type II collagen from mature IVD

A post-translational modification that may have an influence on type IIA/IIB collagen molecular assembly and fibrillogenesis is the 3-hydroxylation of specific proline residues in α1(II) chains [37, 51, 52]. Figure 6 shows mass spectrometry profiles of the α1(II) tryptic peptide containing 3Hyp at P944 from IVD and other type II collagen containing tissues. A tissue specific pattern was observed for 3Hyp occupancy at this site. In the α1(II) collagen chains from bovine vitreous humor, fetal bovine NP and in adult NP, 3Hyp occupancy was 78%, 42% and 22% respectively (Fig. 6, panels E, G, C). The type II collagen fibrils in these tissues are known to be of very thin diameters (< 20nm) [53, 54]. Cartilage has low 3Hyp occupancy (11% in adult and 5% in fetal) (Fig. 6 panels D, H). Cartilage is known to have thicker type II collagen fibrils (>20nm) when compared to the NP and vitreous [46, 54, 55]. Type II collagen from the adult outer annulus showed a higher 3Hyp occupancy (27%) at this site (panel A) than that found in the inner annulus (9%) (panel B) and adult NP (22%) (panel C). It should be reiterated here that, based on the immunofluorescent staining data, type IIA collagen localized to the outer regions of the inner annulus (Figures 4 and 5) and this region was dissected with the outer annulus for collagen analysis by massspectrometry.

**Figure 6.**
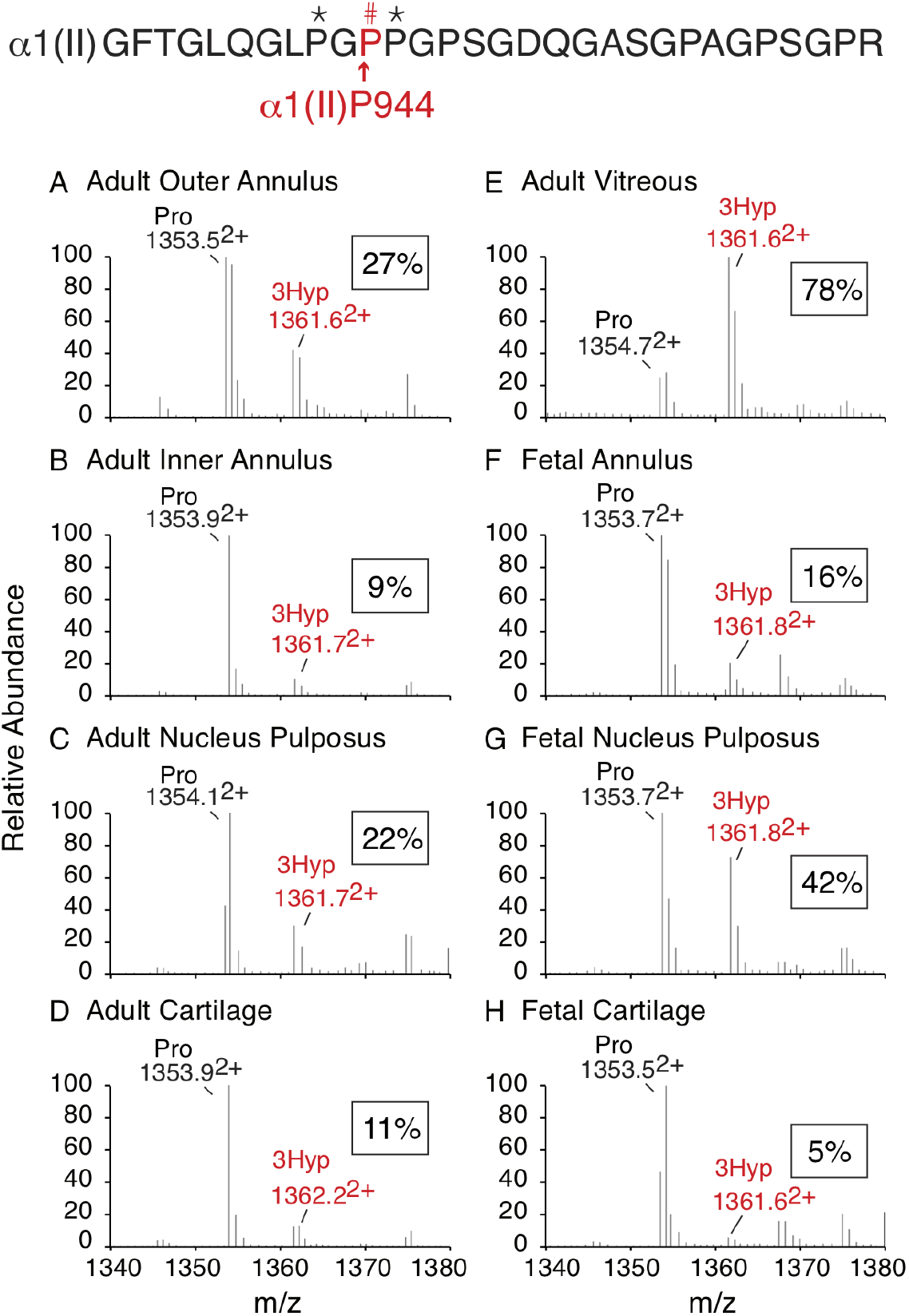
Mass spectrometric profiles of the tryptic peptide containing 3-hydroxyproline at P944 in type II collagen chains. Trypsin digests of type II collagen chains from bovine adult outer annulus fibrosis (A), adult inner annulus fibrosis (B), adult nucleus pulposus (C), adult articular cartilage (D), adult vitreous humor (E), fetal annulus fibrosus, (F), fetal nucleus pulposus (G), fetal epiphyseal cartilage from facet joint in spine (H) were analyzed by mass spectrometry (MS). The peptide of interest is shown at the top of the MS profiles of the tissues. Results for % 3-hydroxyproline (3Hyp) occupancy are shown in boxes.

We also assessed if there were variations in 3Hyp at two other known sites in type II collagen within these tissues. 3Hyp occupancies at three known sites for the various tissues analyzed are summarized in Table 1. P944 and P707 residues are specifically hydroxylated by the prolyl 3-hydroxylase 2 enzyme [50] and we had observed 15-18% 3Hyp at P707 in a week old mouse rib cartilage [36] but no 3Hyp in human NP [52]. Tissue specific variations in 3Hyp occupancy were only observed at the P944 residue in type II collagen. P707 is not 3-hydroxylated in any of the bovine tissues examined here. P986 residue which known to be hydroxylated by the prolyl 3-hydroxylase 1 enzyme [56] is nearly fully 3-hydroxylated in the fetal tissues (90-94%) and adult tissues (80-86%).

**Table 1.** Comparison of 3-Hydroxyproline occupancy in type II collagens from bovine adult and fetal annulus fibrosus (AF), nucleus pulposus (NP), vitreous humor and cartilage. Percentage of 3-Hydroxyproline at each substrate site is shown. The percentages were determined based on the ratio of the m/z peaks for each post-translational variant.

|  | % 3-Hydroxyproline |  |  |
| --- | --- | --- | --- |
|  | P944 | P707 | P986 |
| Adult outer AF | 27 | 0 | 85 |
| Adult inner AF | 14 | 0 | 86 |
| Adult NP | 22 | 0 | 80 |
| Adult cartilage | 11 | 0 | 83 |
| Adult vitreous | 78 | 0 | 86 |
| Fetal AF (inner + outer) | 18 | 0 | 94 |
| Fetal NP | 44 | 0 | 90 |
| Fetal cartilage | 5 | 0 | 94 |

Average of 2 estimates from two different discs and cartilages

## DISCUSSION

Type IIA procollagen protein was immuno-localized in human degenerated intervertebral disc tissue [35]. However, because the human tissue obtained after discectomy was heterogenous in nature, it was not possible to unambiguously determine if the staining was present in regions of IVD tissue or associated the cartilaginous endplate. We were able to biochemically and immunohistochemically address type IIA procollagen expression following careful dissection of bovine intervertebral disc. Western blot analysis clearly shows that intact α1(IIA) procollagen chains (N-propeptide containing the IIA exon 2-encoded cysteine-rich domain + triple helical region) were present in mature steer and calf AF and is absent in NP where only type IIB collagen chains were observed (Figure 2, 3). This finding of IIA procollagen protein expression in normal adult IVD is novel. Type IIA procollagen has been localized to osteoarthritic tissues before [34, 57] as well as in cartilage repair tissue after autologous implantation [58]. We have characterized type IIA procollagen from epiphyseal and rib cartilage from newborn and growing transgenic mice exclusively expressing α1(IIA) isoform [37]. We have also shown that the type IIA procollagen can form heteropolymers with type II, IX and XI in these mice [36].

Immunohistochemistry localized the intact α1(IIA) procollagen chains to the lamellae of the outer margins of the adult inner annulus (Figure 4), a fibro-cartilagenous region normally abundant in both type II and type I collagens [6]. We have observed a similar immunolocalization pattern of type IIA procollagen and type I collagen in human fetal IVD [33]. Here, an intense type IIA procollagen staining was seen in the junction of the inner and outer AF where it co-localized with type I collagen. The exon 2 in α1(IIA) N-propeptide bears similarity to the α1(I) N-propeptide of type I collagen including conserved cysteine residues [29]. From our findings we speculate that the hybrid structure of the α1(IIA) procollagen chains could allow for heteropolymeric assembly of type I and type IIA procollagen in fibrils and confer unique structural properties within fibro-cartilagenous interfaces. Formation of cross-linked types II-type III, V/XI collagen heteropolymers in articular cartilage, NP of the intervertebral disc and meniscus, is not uncommon [59–61] but even in these tissues, what unique structural characteristics such heteropolymers confer is unknown and like for the IVD, necessitates further investigations. It is also important to consider that the α3(XI) chain of type XI procollagen and the α1(II) chain of type II procollagen are encoded by the same gene [62, 63] but the α3(XI) chain is post-translationally over-glycosylated and retains the amino propeptide in the native molecule [64, 65]. We have shown that in newborn and growing transgenic mice exclusively expressing α1(IIA) isoform, α(IIA) procollagen chains can be incorporated into the type XI collagen molecule [36]. So, here in the adult AF, a proportion of molecules having a chain composition of α1(XI) α2(XI) proα1(IIA) could also exist but remains to be validated.

The presence of intact type IIA procollagen in the lamellae of adult inner AF was a surprise since immunoreactivity to type IIA N-propeptide is lost during development [33]. As fetal type II collagen containing tissue develops, exon 2 in the N-propeptide is spliced out [30, 33, 37] and/or the N-propeptide of type IIA/B collagen is processed by ADAMTS-2/3 [66] and although thought to be rapidly degraded by metalloproteinase or internalized by cells [67, 68], we have shown type XI collagen molecules with a chain composition of α1(XI) α2(XI) proα1(IIA) can exist [36]. Interestingly, the C-propeptide trimer of type II collagen has been shown to be a prominent matrix component of immature cartilages and intervertebral disc tissue [42]. Whether the type IIA procollagen detected in the adult AF is part of an existing network of type II/type I collagen fibrils and is somehow exposed or is newly synthesized type IIA procollagen is not clear from our data and will need further investigation. The presence of type IIA procollagen staining in the adult outer annulus is also interesting. Our previous studies showed that the superficial layer of cartilage from the hind limbs of one month-old wild type mice also showed positive staining with the IIA-specific antibody, but it was not clear if this was cleaved type IIA N-propeptide or a proportion of type IIA procollagen chains incorporated in type XI collagen [37]. As expected from the biochemical data (Figures 2, 3), no type IIA propeptide immuno-localized to the NP (Figure 4, D) where the type II collagen triple helix was abundant (Figure 4, H). We have previously shown that triple helical type IIA procollagen is present in the NP of 72 day old developing human disc but by day 101 the type IIA N-propeptide was removed and not detected here after development of the disc. [33]. So, we presume here that type IIA procollagen is either not synthesized in adult NP or if synthesized the N-propeptide was cleaved, rapidly degraded and the epitope lost [67, 68].

It is of note that there are biostructures in the AF that have not yet been biochemically characterized. By high-magnification scanning electron microscopy, Iatridis and ap Gwynn have reported on the presence of distinct small diameter fibrils and collagenous tubules that run parallel to collagen fibers in rat AF [27]. This microtubular network was observed only in sagittal sections of the IVD and not in transverse sections. Interestingly, we detected a robust type IIA procollagen signal in sagittal sections of the inner AF (Figure 4, 5) but in transverse sections faint to no signal was detected (data not shown). Radial cross-bridging elements in AF of sheep discs, bovine tails and adult human discs containing fibrillin have been reported within the AF lamellae [22, 69]. These elements seem to be a consequence of vascular regression during development and stain positive for PECAM-1 an endothelial cell marker [23]. Type IIA procollagen transcript has been localized in vascular tissues as in the epimyocardium of the ventricle and atrium in embryonic mouse heart [32, 70]. Based on the localization of type IIA collagen described here, it is tempting to speculate that fibrils of type IIA procollagen collagen could assemble within these biostructures and play a functional role in this location.

The localization of a proportion of triple helical type II collagen exclusively within the interlamellar regions of the inner annulus (Figure 4, F & J, orange arrows) is also an interesting finding. Type I collagen although normally abundant within the lamellae of the inner and outer AF [6, 7] did not co-localize here (data not shown). It is possible that type II collagen fibrils could confer slightly different mechanical properties to the interlamellar regions of the AF. It is known that under shear, sliding does not occur between AF lamellae and that rigid interlamellar connections are governed primarily by collagen and fibrillin [25]. It is also noteworthy that type II collagen has been observed in regions of tendons that are heavily loaded and also in the enthesis where the extracellular matrix contributes to the ability to withstand compression [71].

What type of fibrils would type IIA collagen be expected to assemble within the fibro-cartilaginous AF? Type IIA collagen has been shown to be associated with thin fibrils in cartilage [46] and the vitreous humor of the eye [54]. We looked for clues in an unique post-translational modification in the α1(II) collagen chain that influences type II collagen fibril assembly and was also elevated in the vitreous humor of the eye [50, 52]. Regions of AF closer to the outer annulus where type IIA procollagen was localized (Figure 4, 5) had a relatively elevated 3Hyp occupancy at P944 (27%) when compared to the type II collagen in the inner annulus and had comparable levels to adult NP (Figure 6, Table 1). In tissues with very thin type II collagen (≤10nm) as in vitreous humor and fetal NP [53, 54], a high occupancy (78%, 42%) was found (Table 1) but in thicker fibrils (≥20nm) of adult and fetal cartilage [46, 54, 55], a low occupancy of 3Hyp at P944 in α1(II) chains (11%, 5%) was observed. In type II collagen from meniscus, another fibrocartilaginous tissue with a decreasing gradient of type II collagen from the inner to the outer zone, a relatively high occupancy of 3Hyp at P944 was also observed [52]. Our data suggests that type IIA collagen in the AF could assemble into thin fibrils. Such smaller diameter fibrils and a tubular network running parallel to the large diameter collagen fibers characteristic of type I collagen was observed by scanning electron microscopy and were assumed to be collagen based. [27]. A prospective more detailed examination of the micro and nano-biostructure by immune-electron microscopy may be able to shed more light on our present observations.

In summary, we biochemically and immunohistochemistry show that type IIA procollagen is present in the AF of mature steer and calf intervertebral disc. Mass spectrometry of type II collagen chains extracted from this region of the AF revealed an elevated level of a specific post translational modification that is normally observed in tissues where thin fibrils of type II collagen assemble. The localization of a proportion of type II collagen exclusively within the inter-lamellae regions of the inner annulus is also an interesting finding. These observations warrant further investigations into understanding intermolecular interactions that help stabilize inter-lamellar adhesion within the intervertebral disc.

## ACKNOWLEDGEMENTS

This work was supported in whole or in part by NIH grants AR 052896 (RJF), AR057025 (RJF), AR053513 (AM), AR036794 (David R. Eyre). We would like to thank, David R Eyre, Jian-Jiu Wu for helpful discussions and Geoffrey Traeger for technical expertise. RJF would like to dedicate this manuscript to the memory of Dr. P. Vijay, a consummate academic, educator and friend.

## Notes

### Competing Interest Statement

The authors have declared no competing interest.

## REFERENCES

[1] A. Nachemson, The Influence of Spinal Movements on the Lumbar Intradiscal Pressure and on the Tensil Stresses in the Annulus Fibrosus, Acta Orthop Scand 33 (1963) 183–207.

[2] S. Gracovetsky, Function of the spine, J Biomed Eng 8(3) (1986) 217–23.

[3] L.A. Setton, J. Chen, Mechanobiology of the intervertebral disc and relevance to disc degeneration, J Bone Joint Surg Am 88 Suppl 2 (2006) 52–7.

[4] S. Roberts, J. Menage, J.P. Urban, Biochemical and structural properties of the cartilage end-plate and its relation to the intervertebral disc, Spine (Phila Pa 1976) 14(2) (1989) 166–74.

[5] J.P. Urban, C.P. Winlove, Pathophysiology of the intervertebral disc and the challenges for MRI, J Magn Reson Imaging 25(2) (2007) 419–32.

[6] D.R. Eyre, H. Muir, Types I and II collagens in intervertebral disc. Interchanging radial distributions in annulus fibrosus, Biochem J 157(1) (1976) 267–70.

[7] D.R. Eyre, H. Muir, Quantitative analysis of types I and II collagens in human intervertebral discs at various ages, Biochim Biophys Acta 492(1) (1977) 29–42.

[8] W.E. Johnson, B. Caterson, S.M. Eisenstein, S. Roberts, Human intervertebral disc aggrecan inhibits endothelial cell adhesion and cell migration in vitro, Spine (Phila Pa 1976) 30(10) (2005) 1139–47.

[9] B. Johnstone, M.T. Bayliss, The large proteoglycans of the human intervertebral disc. Changes in their biosynthesis and structure with age, topography, and pathology, Spine (Phila Pa 1976) 20(6) (1995) 674–84.

[10] J.P. Urban, J.F. McMullin, Swelling pressure of the inervertebral disc: influence of proteoglycan and collagen contents, Biorheology 22(2) (1985) 145–57.

[11] J.P. Urban, A. Maroudas, Swelling of the intervertebral disc in vitro, Connect Tissue Res 9(1) (1981) 1–10.

[12] P.J. Roughley, M. Alini, J. Antoniou, The role of proteoglycans in aging, degeneration and repair of the intervertebral disc, Biochem Soc Trans 30(Pt 6) (2002) 869–74.

[13] A.J. Hayes, M.D. Isaacs, C. Hughes, B. Caterson, J.R. Ralphs, Collagen fibrillogenesis in the development of the annulus fibrosus of the intervertebral disc, Eur Cell Mater 22 (2011) 226–41.

[14] E. Kaapa, X. Han, S. Holm, J. Peltonen, T. Takala, H. Vanharanta, Collagen synthesis and types I, III, IV, and VI collagens in an animal model of disc degeneration, Spine (Phila Pa 1976) 20(1) (1995) 59–66; discussion 66-7.

[15] J.J. Wu, D.R. Eyre, Intervertebral disc collagen. Usage of the short form of the alpha1(IX) chain in bovine nucleus pulposus, J Biol Chem 278(27) (2003) 24521–5.

[16] J.J. Wu, D.R. Eyre, H.S. Slayter, Type VI collagen of the intervertebral disc. Biochemical and electron-microscopic characterization of the native protein, Biochem J 248(2) (1987) 373–81.

[17] S.M. Smith, J. Melrose, Type XI collagen-perlecan-HS interactions stabilise the pericellular matrix of annulus fibrosus cells and chondrocytes providing matrix stabilisation and homeostasis, J Mol Histol 50(3) (2019) 285–294.

[18] C. Berthet-Colominas, A. Miller, D. Herbage, M.C. Ronziere, D. Tocchetti, Structural studies of collagen fibres from intervertebral disc, Biochim Biophys Acta 706(1) (1982) 50–64.

[19] A. Aszodi, D. Chan, E. Hunziker, J.F. Bateman, R. Fassler, Collagen II is essential for the removal of the notochord and the formation of intervertebral discs, J Cell Biol 143(5) (1998) 1399–412.

[20] J. Yu, P.C. Winlove, S. Roberts, J.P. Urban, Elastic fibre organization in the intervertebral discs of the bovine tail, J Anat 201(6) (2002) 465–75.

[21] J. Yu, J.C. Fairbank, S. Roberts, J.P. Urban, The elastic fiber network of the anulus fibrosus of the normal and scoliotic human intervertebral disc, Spine (Phila Pa 1976) 30(16) (2005) 1815–20.

[22] J. Yu, U. Tirlapur, J. Fairbank, P. Handford, S. Roberts, C.P. Winlove, Z. Cui, J. Urban, Microfibrils, elastin fibres and collagen fibres in the human intervertebral disc and bovine tail disc, J Anat 210(4) (2007) 460–71.

[23] L.J. Smith, D.M. Elliott, Formation of lamellar cross bridges in the annulus fibrosus of the intervertebral disc is a consequence of vascular regression, Matrix Biol 30(4) (2011) 267–74.

[24] J. Tavakoli, D.M. Elliott, J.J. Costi, Structure and mechanical function of the interlamellar matrix of the annulus fibrosus in the disc, J Orthop Res 34(8) (2016) 1307–15.

[25] A.J. Michalek, M.R. Buckley, L.J. Bonassar, I. Cohen, J.C. Iatridis, Measurement of local strains in intervertebral disc anulus fibrosus tissue under dynamic shear: contributions of matrix fiber orientation and elastin content, J Biomech 42(14) (2009) 2279–85.

[26] J. Yu, M.L. Schollum, K.R. Wade, N.D. Broom, J.P. Urban, ISSLS Prize Winner: A Detailed Examination of the Elastic Network Leads to a New Understanding of Annulus Fibrosus Organization, Spine (Phila Pa 1976) 40(15) (2015) 1149–57.

[27] J.C. Iatridis, I. ap Gwynn, Mechanisms for mechanical damage in the intervertebral disc annulus fibrosus, J Biomech 37(8) (2004) 1165–75.

[28] I. ap Gwynn, S. Wade, M.J. Kaab, G.R. Owen, R.G. Richards, Freeze-substitution of rabbit tibial articular cartilage reveals that radial zone collagen fibres are tubules, J Microsc 197(Pt 2) (2000) 159–72.

[29] M.C. Ryan, L.J. Sandell, Differential expression of a cysteine-rich domain in the amino-terminal propeptide of type II (cartilage) procollagen by alternative splicing of mRNA, J Biol Chem 265(18) (1990) 10334–9.

[30] L.J. Sandell, A.M. Nalin, R.A. Reife, Alternative splice form of type II procollagen mRNA (IIA) is predominant in skeletal precursors and non-cartilaginous tissues during early mouse development, Dev Dyn 199(2) (1994) 129–40.

[31] A. Oganesian, Y. Zhu, L.J. Sandell, Type IIA procollagen amino propeptide is localized in human embryonic tissues, J Histochem Cytochem 45(11) (1997) 1469–80.

[32] L.J. Ng, P.P. Tam, K.S. Cheah, Preferential expression of alternatively spliced mRNAs encoding type II procollagen with a cysteine-rich amino-propeptide in differentiating cartilage and nonchondrogenic tissues during early mouse development, Dev Biol 159(2) (1993) 403–17.

[33] Y. Zhu, A. McAlinden, L.J. Sandell, Type IIA procollagen in development of the human intervertebral disc: regulated expression of the NH(2)-propeptide by enzymic processing reveals a unique developmental pathway, Dev Dyn 220(4) (2001) 350–62.

[34] T. Aigner, Y. Zhu, H.H. Chansky, F.A. Matsen, 3rd, W.J. Maloney, L.J. Sandell, Reexpression of type IIA procollagen by adult articular chondrocytes in osteoarthritic cartilage, Arthritis Rheum 42(7) (1999) 1443–50.

[35] Y. Zhu, J.J. Wu, M.A. Weis, S.K. Mirza, D.R. Eyre, Type IX collagen neo-deposition in degenerative discs of surgical patients whether genotyped plus or minus for COL9 risk alleles, Spine (Phila Pa 1976) 36(24) (2011) 2031–8.

[36] A. McAlinden, G. Traeger, U. Hansen, M.A. Weis, S. Ravindran, L. Wirthlin, D.R. Eyre, R.J. Fernandes, Molecular properties and fibril ultrastructure of types II and XI collagens in cartilage of mice expressing exclusively the alpha1(IIA) collagen isoform, Matrix Biol 34 (2014) 105–13.

[37] R. Lewis, S. Ravindran, L. Wirthlin, G. Traeger, R.J. Fernandes, A. McAlinden, Disruption of the developmentally-regulated Col2a1 pre-mRNA alternative splicing switch in a transgenic knock-in mouse model, Matrix Biol 31(3) (2012) 214–26.

[38] R.J. Fernandes, D.J. Wilkin, M.A. Weis, W.R. Wilcox, D.H. Cohn, D.L. Rimoin, D.R. Eyre, Incorporation of structurally defective type II collagen into cartilage matrix in kniest chondrodysplasia, Arch Biochem Biophys 355(2) (1998) 282–90.

[39] R.J. Fernandes, R.E. Seegmiller, W.R. Nelson, D.R. Eyre, Protein consequences of the Col2a1 C-propeptide mutation in the chondrodysplastic Dmm mouse, Matrix Biol 22(5) (2003) 449–53.

[40] R.J. Fernandes, M. Weis, M.A. Scott, R.E. Seegmiller, D.R. Eyre, Collagen XI chain misassembly in cartilage of the chondrodysplasia (cho) mouse, Matrix Biol 26(8) (2007) 597–603.

[41] R.J. Fernandes, T.M. Schmid, M.A. Harkey, D.R. Eyre, Incomplete processing of type II procollagen by a rat chondrosarcoma cell line, Eur J Biochem 247(2) (1997) 620–4.

[42] C. Niyibizi, J.J. Wu, D.R. Eyre, The carboxypropeptide trimer of type II collagen is a prominent component of immature cartilages and intervertebral-disc tissue, Biochim Biophys Acta 916(3) (1987) 493–9.

[43] R.J. Fernandes, T.M. Schmid, D.R. Eyre, Assembly of collagen types II, IX and XI into nascent hetero-fibrils by a rat chondrocyte cell line, Eur J Biochem 270(15) (2003) 3243–50.

[44] G.A. Whitney, T.J. Kean, R.J. Fernandes, S. Waldman, M.Y. Tse, S.C. Pang, J.M. Mansour, J.E. Dennis, Thyroxine Increases Collagen Type II Expression and Accumulation in Scaffold-Free Tissue-Engineered Articular Cartilage, Tissue Eng Part A 24(5-6) (2018) 369–381.

[45] J.E. Dennis, G.A. Whitney, J. Rai, R.J. Fernandes, T.J. Kean, Physioxia Stimulates Extracellular Matrix Deposition and Increases Mechanical Properties of Human Chondrocyte-Derived Tissue-Engineered Cartilage, Front Bioeng Biotechnol 8 (2020) 590743.

[46] Y. Zhu, A. Oganesian, D.R. Keene, L.J. Sandell, Type IIA procollagen containing the cysteine-rich amino propeptide is deposited in the extracellular matrix of prechondrogenic tissue and binds to TGF-beta1 and BMP-2, J Cell Biol 144(5) (1999) 1069–80.

[47] M.A. Cremer, A.H. Kang, Collagen-induced arthritis in rodents: a review of immunity to type II collagen with emphasis on the importance of molecular conformation and structure, Int Rev Immunol 4(1) (1988) 65–81.

[48] R.J. Fernandes, M. Weis, M.A. Scott, R.E. Seegmiller, D.R. Eyre, Collagen XI chain misassembly in cartilage of the chondrodysplasia (cho) mouse, Matrix Biol 26(8) (2007) 597–603.

[49] R.J. Fernandes, M.A. Harkey, M. Weis, J.W. Askew, D.R. Eyre, The post-translational phenotype of collagen synthesized by SAOS-2 osteosarcoma cells, Bone 40(5) (2007) 1343–51.

[50] R.J. Fernandes, A.W. Farnand, G.R. Traeger, M.A. Weis, D.R. Eyre, A role for prolyl 3-hydroxylase 2 in post-translational modification of fibril-forming collagens, J Biol Chem 286(35) (2011) 30662–9.

[51] A.D. Murdoch, T.E. Hardingham, D.R. Eyre, R.J. Fernandes, The development of a mature collagen network in cartilage from human bone marrow stem cells in Transwell culture, Matrix Biol 50 (2016) 16–26.

[52] M.A. Weis, D.M. Hudson, L. Kim, M. Scott, J.J. Wu, D.R. Eyre, Location of 3-hydroxyproline residues in collagen types I, II, III, and V/XI implies a role in fibril supramolecular assembly, J Biol Chem 285(4) (2010) 2580–90.

[53] H. Inoue, T. Takeda, Three-dimensional observation of collagen framework of lumbar intervertebral discs, Acta Orthop Scand 46(6) (1975) 949–56.

[54] A. Reardon, L. Sandell, C.J. Jones, D. McLeod, P.N. Bishop, Localization of pN-type IIA procollagen on adult bovine vitreous collagen fibrils, Matrix Biol 19(2) (2000) 169–73.

[55] P.N. Bishop, D.F. Holmes, K.E. Kadler, D. McLeod, K.J. Bos, Age-related changes on the surface of vitreous collagen fibrils, Invest Ophthalmol Vis Sci 45(4) (2004) 1041–6.

[56] R. Morello, T.K. Bertin, Y. Chen, J. Hicks, L. Tonachini, M. Monticone, P. Castagnola, F. Rauch, F.H. Glorieux, J. Vranka, H.P. Bachinger, J.M. Pace, U. Schwarze, P.H. Byers, M. Weis, R.J. Fernandes, D.R. Eyre, Z. Yao, B.F. Boyce, B. Lee, CRTAP is required for prolyl 3-hydroxylation and mutations cause recessive osteogenesis imperfecta, Cell 127(2) (2006) 291–304.

[57] T.P. Lefkoe, A.M. Nalin, J.M. Clark, R.A. Reife, J. Sugai, L.J. Sandell, Gene expression of collagen types IIA and IX correlates with ultrastructural events in early osteoarthrosis: new applications of the rabbit meniscectomy model, J Rheumatol 24(6) (1997) 1155–63.

[58] S. Roberts, J. Menage, L.J. Sandell, E.H. Evans, J.B. Richardson, Immunohistochemical study of collagen types I and II and procollagen IIA in human cartilage repair tissue following autologous chondrocyte implantation, Knee 16(5) (2009) 398–404.

[59] D.R. Eyre, J.J. Wu, Collagen of fibrocartilage: a distinctive molecular phenotype in bovine meniscus, FEBS Lett 158(2) (1983) 265–70.

[60] J.J. Wu, M.A. Weis, L.S. Kim, B.G. Carter, D.R. Eyre, Differences in chain usage and cross-linking specificities of cartilage type V/XI collagen isoforms with age and tissue, J Biol Chem 284(9) (2009) 5539–45.

[61] J.J. Wu, M.A. Weis, L.S. Kim, D.R. Eyre, Type III collagen, a fibril network modifier in articular cartilage, J Biol Chem 285(24) (2010) 18537–44.

[62] R.E. Burgeson, D.W. Hollister, Collagen heterogeneity in human cartilage: identification of several new collagen chains, Biochem Biophys Res Commun 87(4) (1979) 1124–31.

[63] C.A. Reese, R. Mayne, Minor collagens of chicken hyaline cartilage, Biochemistry 20(19) (1981) 5443–8.

[64] D.R. Eyre, J.J. Wu, Type XI or 1a2a3a collagen, in: R. Mayne, Burgeson, RE (Ed.), Structure and Function of Collagen Types, Academic Press, New York, 1987, pp. 261–281.

[65] J.R. Thom, N.P. Morris, Biosynthesis and proteolytic processing of type XI collagen in embryonic chick sterna, J Biol Chem 266(11) (1991) 7262–9.

[66] R.J. Fernandes, S. Hirohata, J.M. Engle, A. Colige, D.H. Cohn, D.R. Eyre, S.S. Apte, Procollagen II amino propeptide processing by ADAMTS-3. Insights on dermatosparaxis, J Biol Chem 276(34) (2001) 31502–9.

[67] A. McAlinden, Y. Zhu, L.J. Sandell, Expression of type II procollagens during development of the human intervertebral disc, Biochem Soc Trans 30(Pt 6) (2002) 831–8.

[68] N. Fukui, A. McAlinden, Y. Zhu, E. Crouch, T.J. Broekelmann, R.P. Mecham, L.J. Sandell, Processing of type II procollagen amino propeptide by matrix metalloproteinases, J Biol Chem 277(3) (2002) 2193–201.

[69] C.A. Pezowicz, P.A. Robertson, N.D. Broom, The structural basis of interlamellar cohesion in the intervertebral disc wall, J Anat 208(3) (2006) 317–30.

[70] V.C. Lui, L.J. Ng, J. Nicholls, P.P. Tam, K.S. Cheah, Tissue-specific and differential expression of alternatively spliced alpha 1(II) collagen mRNAs in early human embryos, Dev Dyn 203(2) (1995) 198–211.

[71] M. Benjamin, J.R. Ralphs, Fibrocartilage in tendons and ligaments--an adaptation to compressive load, J Anat 193 (Pt 4) (1998) 481–94.

